# A transposon origin of large tandem repeats in the butterfly *Papilio bianor*

**DOI:** 10.64898/2026.07.30.741882

**Authors:** Tianzhu Xiong, Daniel A. Barbash

## Abstract

Without single centromeres, holocentric chromosomes do not accumulate repetitive DNA in the same way as their monocentric ancestors. In holocentric butterflies and moths, large tandem repeats are rare compared to other insects with monocentric chromosomes. We discover in the butterfly *Papilio bianor* that nearly half of its chromosomes have large tandem repeats composed entirely of transposons. These transposon arrays are formed by concerted expansion of multiple families, often locked together as a single repeating unit of 100kb-200kb in length. On Hi-C contact maps, transposon arrays partition chromosomes into distinct “arms” and insulate the inter-arm contact. Chromosomes with transposon arrays are also more compact than those without the arrays. In male meiosis, transposon arrays facilitate double crossovers, likely by reducing crossover interference between the flanking arms. As a result, chromosomes with transposon arrays exhibit increased recombination rates. Taken together, we demonstrate that large tandem repeats can affect multiple aspects of holocentric chromosomes, and in many ways, the effects resemble those of centromeres in monocentric organisms.

## 1. Introduction

In monocentric organisms, centromeres attract kinetochores during cell division, and they often harbor large blocks of repeats composed of satellites and transposons [1–3]. These centromeric repeats have prominent effects on chromosomes. For instance, constitutive heterochromatin encompasses centromeric repeats and suppresses their expression [4,5]. This architecture also frequently results in different centromeres clustering together in interphase nuclei [6,7]. In meiosis, crossover rarely occurs in centromeric repeats [8]. Centromeric repeats also insulate the spatial contact between flanking sequences in interphase nuclei, thus individualizing chromosome arms [6].

In contrast, holocentric genomes lack single centromeres, which are usually hotspots for repeats [9,10]. Consequently, tandem repeats may accumulate elsewhere on holocentric chromosomes, with effects potentially distinct from those associated with single centromeres. Some holocentric lineages employ multiple repetitive regions across a single chromosome as kinetochore assembly sites [11,12], but this is not the case for the holocentric order Lepidoptera (butterflies and moths). For silk moth *Bombyx mori*, mitotic kinetochores assemble dynamically throughout the chromosomal axis, with a preference for silent chromatin, and it is likely sequence-independent [13]. Coincidentally, Lepidopteran autosomes so far studied lack highly repetitive regions comparable to centromeres [14], and large tandem repeats spanning multiple megabases are particularly rare except for telomeres. Thus, it is unclear how large tandem repeats would affect Lepidopteran chromosomes.

In an existing genome assembly of the butterfly *Papilio bianor*, we identified that nearly half of its chromosomes have gaps at large tandem repeats, many of which are still poorly assembled [15,16]. The abundance of tandem repeats in *P. bianor* provides an exceptional opportunity to study their effects on holocentric chromosomes. We first reassemble its genome to improve contiguity and correct for assembly artifacts. Then, we characterize the structure of tandem repeats and their impact on chromosome properties.

## 2. Results

### (a) Recent tandem expansion of transposons in *Papilio bianor*

We produced a new primary assembly using the published PacBio HiFi reads and Hi-C reads [16]. The new assembly corrected numerous small-scale errors in the old assembly and improved contiguity (**Supplementary File S1-Table S1**). Among the 29 autosomes and the Z chromosome, we discovered that 16 of them resemble the chromosome structure of *B. mori* [17], namely, repeats are distributed across the chromosome and are enriched only at telomeres (**Figure 1A, Type α**). On the contrary, the remaining 14 chromosomes contain long stretches of repetitive DNA enriched for transposons. The majority of tandemly organized transposons are internal to the chromosomes (**Figure 1A, Type β**), whereas some accumulate at chromosome ends (**Figure 1A, Type γ**). We could not fill the gaps present in many transposon arrays despite our extensive optimization of parameters in the assembly software hifiasm [18]. Even with gaps, many arrays already approach 5mb in length, suggesting that their actual sizes could be even greater.

**Figure 1.**
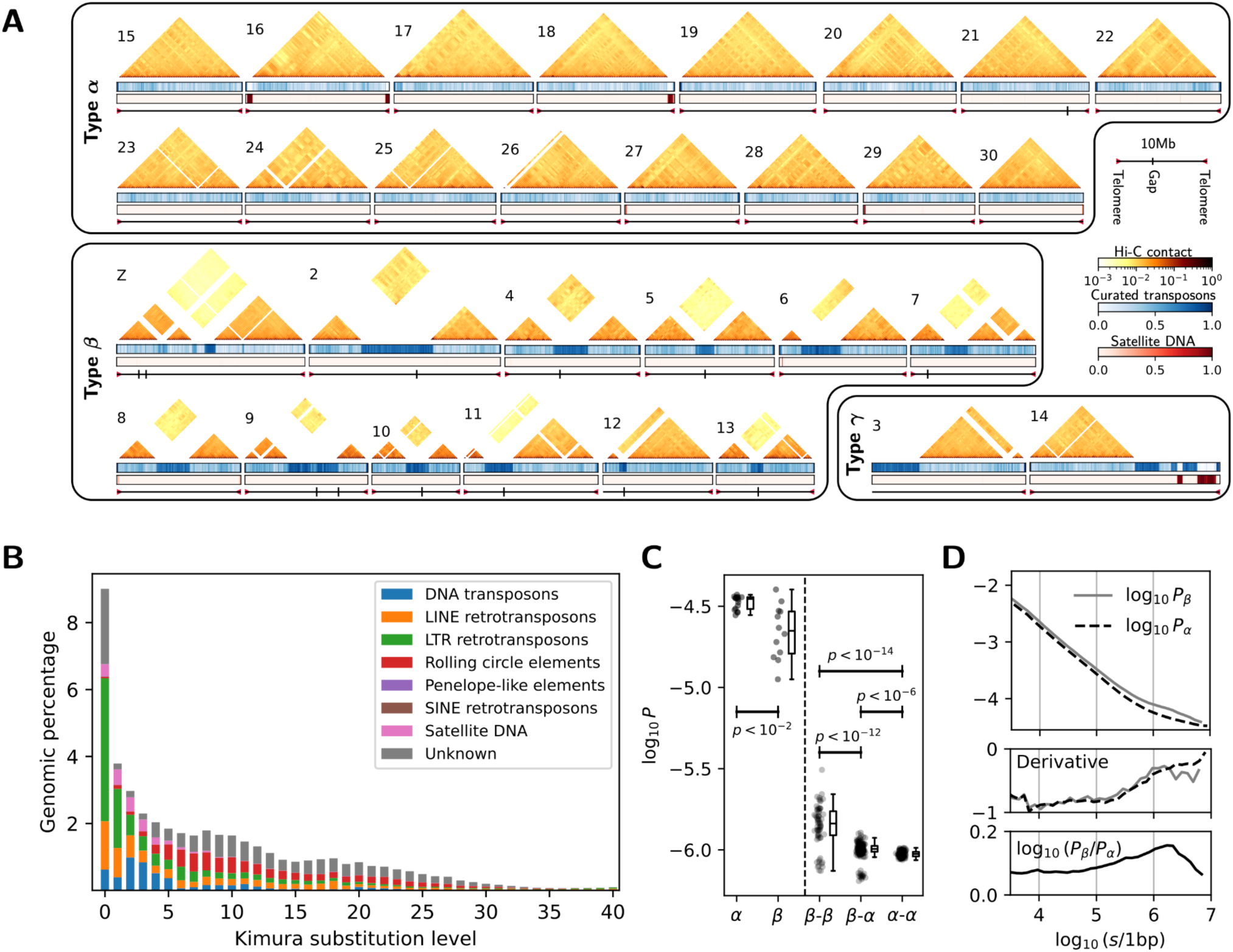
The properties of the new *P. bianor* assembly. **(A)** 30 chromosomes are classified as Type α, β, or γ depending on the presence/locations of transposon (TE) arrays. Orange heatmaps are 250kb-resolution Hi-C contacts, and blank stripes correspond to repetitive regions refractory to Hi-C read mapping. Blue/red tracks are transposon/satellite DNA density per 50kb window. Telomeres and gaps are depicted as small triangles and vertical lines, respectively. The scale bar is shown for a 10mb sequence on the right. **(B)** The Kimura substitution levels for all major classes of repetitive DNA. **(C)** Left: the average *cis*-contact between the terminal 1mb windows for each Type α and β chromosome. Right: the average *trans*-contact between all 1mb windows for each pair of different Type α and β chromosomes. Hi-C matrices are at a resolution of 1kb. P-values are from *t*-tests with unequal variances. **(D)** Top: the *P*(*s*) curves for Type α and β chromosomes. Middle: the log-log derivative of the *P*(*s*) curves. Bottom: The log-ratio of *P*(*s*) curves between the two types of chromosomes. *s* is in the unit of 1bp. *P* is from Hi-C matrices with a resolution of 1kb.

The transposon arrays appear to result from recent tandem expansion (**Figure 1B**). The Kimura substitution level (a metric of sequence divergence) for manually curated transposons annotated genome-wide is skewed strongly to the left, indicating that most copies in the same transposon family are closely related. DNA transposons, LINE, LTR, and several unknown elements all contribute to the recent expansions (**Figure S1**). A notable exception is that satellite DNA contributes little to the repeat landscape, despite its frequent presence in tandem DNA arrays of many other organisms.

### (b) Transposon arrays alter chromosomal contacts

To study whether transposon arrays alter the 3D conformation of interphase chromosomes, we analyzed Hi-C contact frequencies (*P*) for each chromosome (**Figure 1A**). We excluded Type γ chromosomes in this analysis due to the small sample size.

For long-range contacts, Type β chromosomes have internal transposon arrays, and concomitantly, Hi-C contacts between the flanking sequences (“arms”) of transposon arrays are visually weaker than contacts between even the most distant loci on Type α chromosomes. On the other hand, contacts within each “arm” appear to remain intact. Consistently, *cis*-contacts between the two most distant 1mb windows are weaker and more variable on Type β chromosomes than those on Type α chromosomes (**Figure 1C, left**). In contrast, *trans*-contacts are stronger between different Type β chromosomes, but weaker between different Type α chromosomes (**Figure 1C, right**). The elevated *trans*-contact between Type β chromosomes is not due to stable clustering among transposon arrays, otherwise flanking regions of arrays should display signals of strong contact, which are absent on the global Hi-C map (**Figure S2**). These data indicate that, on average, transposon arrays insulate *cis*-contacts between the flanking “arms” of a single chromosome but also bring different “arms” from different chromosomes closer together.

Among short-range contacts (distance *s<*1mb), Type α and β chromosomes conform to a similar standard *P*(*s*) curve with a log-log derivative close to -1, a widely observed phenomenon across many Eukaryotes (**Figure 1D, top, middle**) [19]. However, the magnitude of contact is slightly higher for Type β chromosomes at this distance scale (**Figure 1D, bottom**), particularly when *s* is close to 1mb. We interpret it as evidence that Type β chromosomes are, on average, more compact than Type α chromosomes at short distances, possibly caused by transposon arrays.

### (c) Higher-order structures in transposon arrays

The high level of similarity among transposon copies prompted us to investigate the structure within transposon arrays. We computed a k-mer-based sequence similarity score between pairs of 100kb windows to visualize higher-order structures among the 14 largest transposon arrays (**Figure 2A**). At this length scale, most arrays are similar, but the transposon array on the Z chromosome is visually an outlier dissimilar to all others. The array on Chromosome 14 contains two blocks of satellite DNA (**Figure 1A**), but otherwise is similar to other arrays.

**Figure 2.**
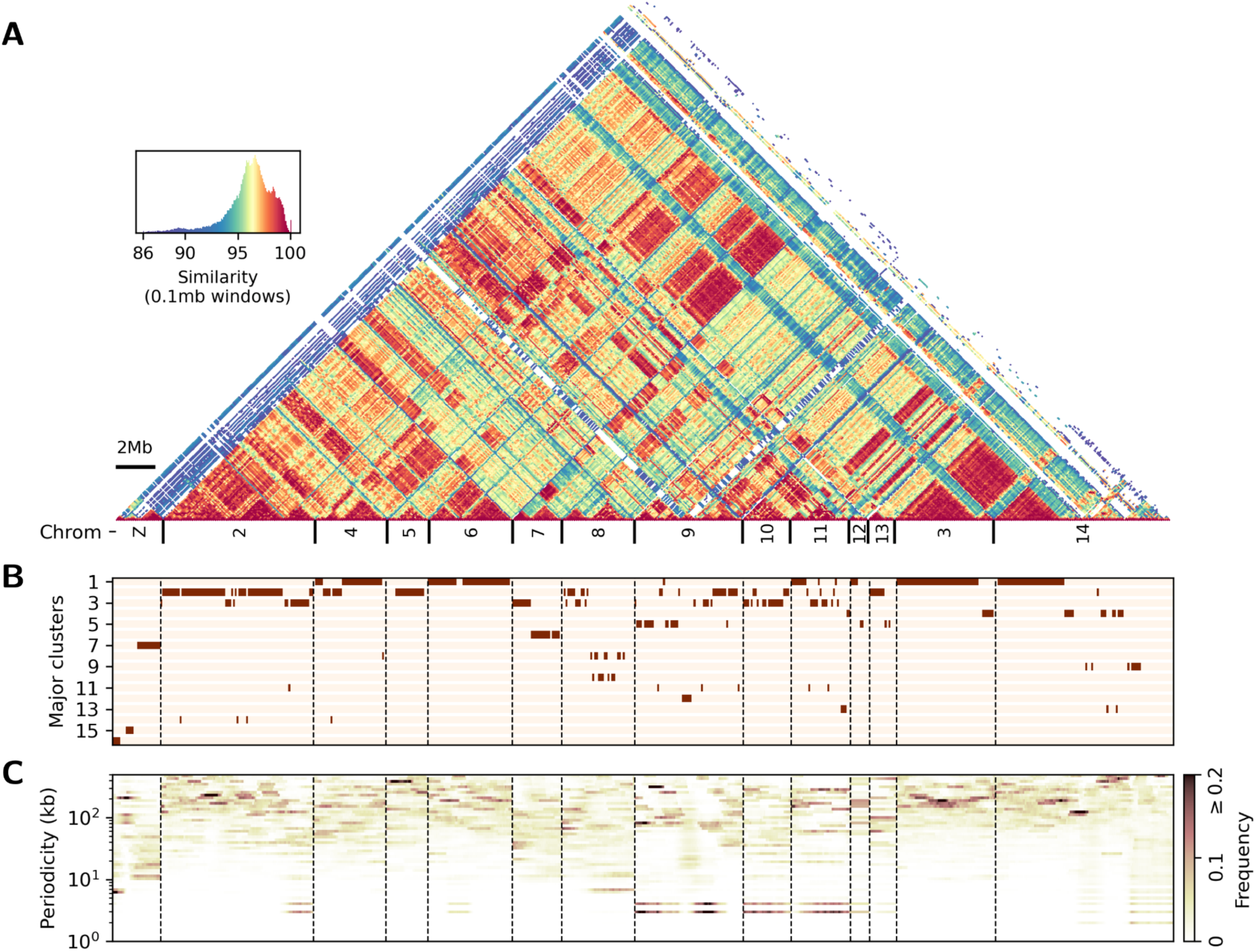
Higher-order structures in the transposon arrays. The x-axes of the three subplots are aligned. **(A)** The heatmap (also known as “dotplot”) of similarity scores between each pair of 0.1mb windows across all 14 major transposon arrays. Numbers in the bottom row indicate the chromosomal origin of each array. **(B)** Major clusters identified via similarity scores at 0.1mb resolution. The top 16 clusters are ranked by their total sizes on the 14 arrays. Additional clusters are not shown. **(C)** Periodicity calculated using a dotplot at 1kb-resolution for consecutive 1mb windows with a step size of 10kb. Each column is interpreted as an independent histogram summarizing the distribution of periodicities in that 1mb window. For instance, most histograms in the Chr3 array peak at periodicities between 100kb-200kb, whereas most histograms in the Chr10 array have a peak periodicity <5kb.

To delineate boundaries between different higher-order structures, we applied a clustering algorithm to similarity scores across all 100kb windows, so that windows within the same cluster are highly alignable to each other and belong to a similar higher-order structure. We display the first 16 clusters ranked by size in **Figure 2B**. Again, it shows that the Z chromosome contains distinct clusters not found in any autosomal arrays. Abundant clusters on autosomes, however, often spread over multiple chromosomes. For instance, the two most abundant clusters (cluster 1 and cluster 2) span a large portion of the arrays on chromosomes 2, 4, 5, 6, 3, and 14, indicating that many autosomal arrays may result from a shared expansion event.

To understand how transposons are organized within each cluster, we zoomed in to a finer scale (1kb windows), and computed sequence similarity scores again between pairs of windows. At this scale, the window size is smaller than the typical size of a transposon, so it is possible to identify pairs of almost identical windows (similarity score > 99). The spacing between adjacent identical windows provides information about the size of the repeating units, which we call “periodicity”. If transposons are organized as higher-order repeats in the array, we expect to find a definitive periodicity matching the size of the higher-order repeating unit, whereas randomly organized transposons should generate more randomly distributed periodicities. We plotted the distribution of periodicity within each 1mb sequence in **Figure 2C**. While some clusters have predominantly small periodicities (e.g., cluster 3), the two most common clusters have periodicities well above 100kb --- much greater than the size of typical transposons (an example dotplot for cluster 1 with ∼200kb periodicity is shown in **Figure S3**).

Although the exact lengths of the repeating units are variable, large periodicities imply that a series of transposons must be grouped together as a single higher-order repeating unit. We plotted the occupancy of each high-level transposon family in representative arrays. For the autosomal clusters 1 & 2 of large periodicities, DNA transposons, LTR and LINE retrotransposons are all abundant (**Figure 3A, 3B**), whereas the autosomal cluster 3 is dominated by unknown elements of a much shorter periodicity (**Figure 3C**). The Z-linked cluster 7 is uniquely enriched for LTR/Pao retrotransposons (**Figure 3D**), whereas LTR/Gypsy retrotransposons are more common in autosomal clusters. Thus, the recent expansions of multiple families of transposons in **Figure 1B** are likely not independent events. Instead, a specific mechanism locks a consecutive series of transposons together, then the entire series expands tandemly as a single unit.

**Figure 3.**
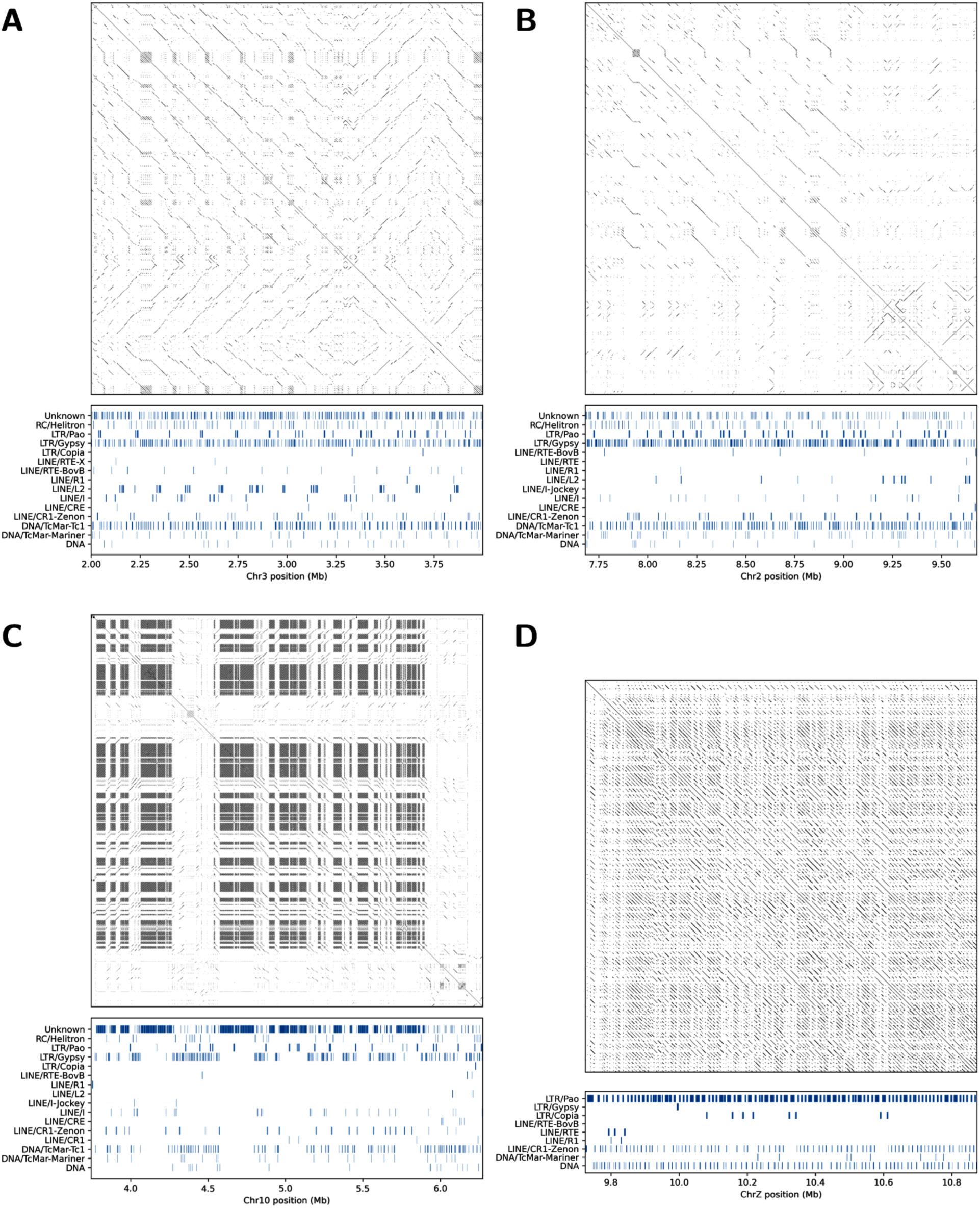
Transposon components of several representative clusters. The resolution for all dotplots is 1kb. **(A)** Chromosome 3, cluster 1. **(B)** Chromosome 2, cluster 2. **(C)** Chromosome 10, cluster 3. **(D)** Chromosome Z, cluster 7.

### (d) Transposon arrays facilitate double crossovers in hybrid males

Since transposons in many organisms affect meiotic crossover patterns [20,21], we sought to study the effect of transposon arrays on paternal crossovers (female meiosis is achiasmatic in Lepidoptera). Using an existing panel of backcross lines sequenced for an interspecific cross between *P. bianor* and *P. dehaanii* (**Figure 4D**) [22], we mapped recombination breakpoints onto our new assembly. First, by counting breakpoints, we estimated that different chromosomes have different frequencies of forming double crossovers in paternal meiosis (**Figure 4A**). Curiously, chromosomes with the most frequent double crossovers appear to be Type β chromosomes, some of which form double crossovers at a rate approaching 50%. This rate is much higher than that of typical butterfly meiosis, which often has almost complete crossover interference with only one crossover per chromosome pair [23].

**Figure 4.**
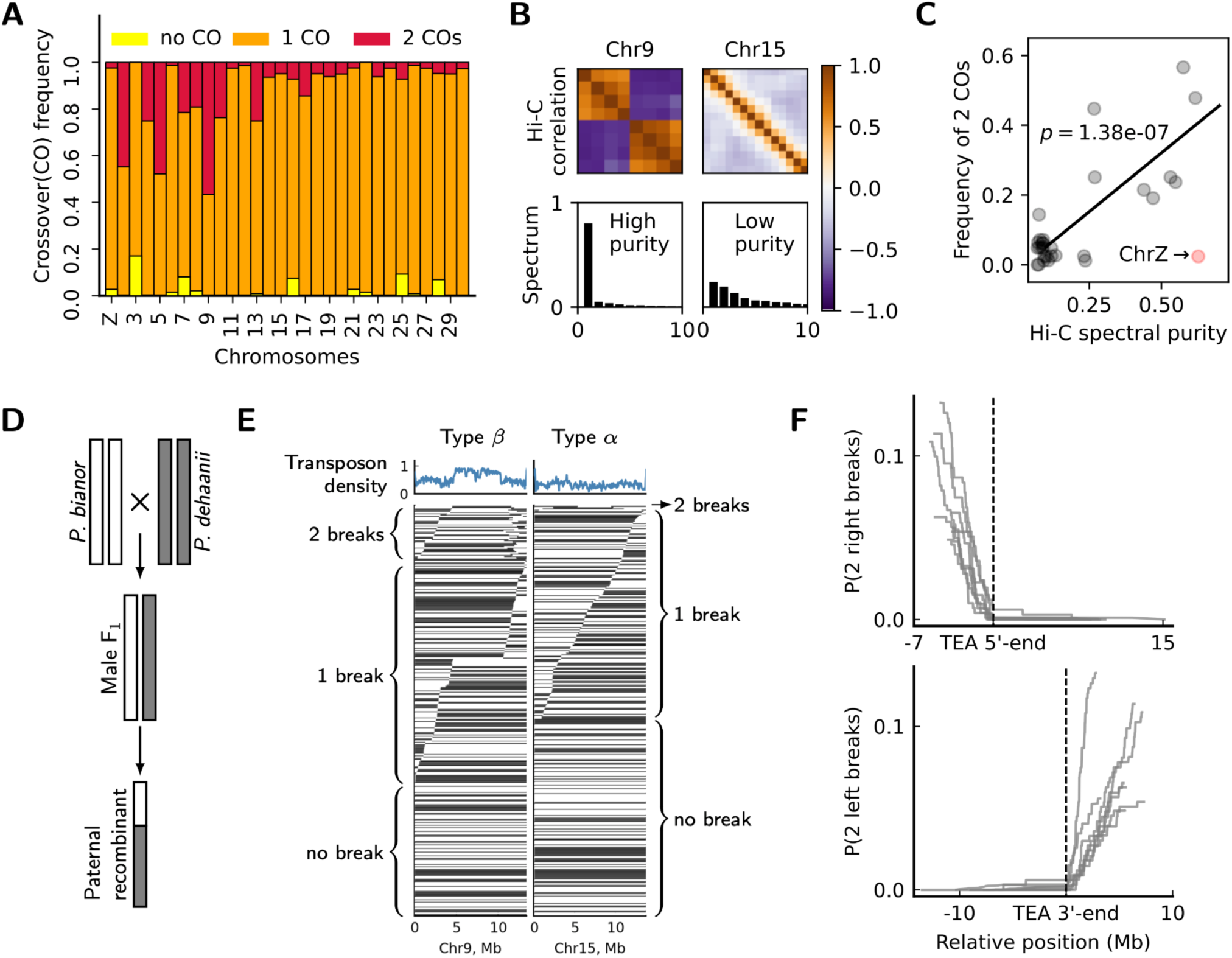
Transposon arrays facilitate double crossovers. **(A)** Estimated frequencies of different numbers of crossovers per chromosome pair per meiosis. Chr15-30 are Type α chromosomes and are among the least likely to have double crossovers. **(B)** Examples of Hi-C correlation matrices and spectral purity between typical Type α (Chr15) and Type β (Chr9) chromosomes. The matrices are at a resolution of 1mb. **(C)** The frequency of double crossovers is positively correlated with a measure of spectral purity. The Z chromosome is visibly an outlier. p-value and regression are computed without the Z chromosome. **(D)** The backcross scheme to infer paternal meiotic crossover patterns. All three generations were sequenced including 335 paternal recombinants. **(E)** Examples of recombinant haplotypes for Type α (Chr15) and Type β (Chr9) chromosomes. The top plots show transposon density on the whole chromosome. In the bottom plots, each horizontal line is a unique recombinant haplotype, and grandparental ancestries are designated as gray and white. The brackets group haplotypes with different numbers of breakpoints together. In each bracket, haplotypes are sorted by the position of the first breakpoint. **(F)** Top: the probability of having two breakpoints to the 3’-side of a given position on the x-axis. The position on the x-axis is relative to the 5’-end of the transposon array (TEA) on each chromosome (dashed line). Bottom: similarly, the probability of having two breakpoints to the 5’-side of a given position on the x-axis. The position on the x-axis is relative to the 3’-end of the transposon array (TEA) on each chromosome (dashed line).

A bipartite Hi-C pattern also coincides with double crossovers. To quantify the degree of Hi-C bipartition, we computed the Hi-C correlation matrix at 1mb resolution for each chromosome and extracted its eigenspectrum (**Figure 4B**). For chromosomes with internal transposon arrays (e.g., chromosome 9), their Hi-C contact matrices are bipartite due to the insulating effects of the arrays. Correspondingly, their Hi-C correlation matrices are also bipartite, and are associated with a concentrated spectrum of higher spectral purity. In contrast, chromosomes without transposon arrays (e.g., chromosome 15) tend to have a flatter spectrum of lower spectral purity. The frequency of double crossovers is positively correlated with Hi-C spectral purity (**Figure 4C**), indicating that the degree of Hi-C bipartition predicts the frequency of double crossovers. Interestingly, the Z chromosome is again an outlier, which is strongly bipartite on the Hi-C plot but has minimal double crossovers.

Finally, we plotted recombination breakpoints from all paternal haplotypes to study the spatial distribution of double crossovers (**Figure 4E**). It appears that if two breakpoints occur on a single haplotype, they almost always flank the internal transposon array (e.g., chr9). Indeed, for the eight chromosomes with the most frequent double crossovers, the probability of *having two breakpoints to the 3’-side of a given position* rises rapidly only when the position moves to the 5’-side of the 5’-end of the transposon array (**Figure 4F, top**). Correspondingly, the probability of *having two breakpoints to the 5’-side of a given position* rises rapidly only when the position moves to the 3’-side of the 3’-end of the transposon array (**Figure 4F, bottom**). In other words, most double breakpoints must flank the entire transposon array. Since double breakpoints are necessarily created by double crossovers, it is likely that most double crossovers must also flank the entire transposon array.

Taken together, our data support a hypothesis in which transposon arrays weaken crossover interference that normally spans the entire chromosome. Thus, double crossovers form readily on the opposite sides of the transposon array. Except for the Z chromosome, most Type β chromosomes have increased overall recombination rates from this effect.

## 3. Discussion

### (a) The origin of transposon arrays

Many recently assembled Lepidopteran genomes from the Darwin’s Tree of Life Project [14] and Project Psyche [24] resemble the typical chromosomes in *B. mori* [17] as well as the Type α chromosomes in *P. bianor*. It is surprising to find abundant tandem repeats in *P. bianor*, because other species from the same genus *Papilio* with published genomes, such as *P. machaon*, *P. polytes*, and *P. xuthus* [25–27], do not possess large transposon arrays (**Figure S4**). Thus, it is likely that such repeats, along with Type β/γ chromosomes, are unique to this or closely related species in the genus *Papilio*.

Another mystery is that so many of the transposon arrays have a periodicity on the order of 100kb. Since this length scale is much greater than the size of a typical transposon, there must be a distinct mechanism to copy/insert the whole series of transposons elsewhere in the array. Such a large repeating unit in tandem DNA arrays has rarely been reported, and certainly does not fall under the typical definition of satellite DNA [28,29]. The abundance of such units also differs from that of segmental duplications, which typically exist as only a handful of copies [30]. We hypothesize that one of the families from the arrays, the rolling circle transposons, might be responsible for the formation of these unusual arrays. Rolling circle transposons, also known as Helitrons, replicate genomic DNA from designated start sites until hitting a stop site defined by a characteristic motif [31]. The replicated segment is then circularized and inserted into a new genomic location. It is plausible that a Helitron stop site might have mutated within an existing transposon array. Without the stop site, rolling circle replication proceeds indefinitely through the transposon array, picking up a large number of different classes of transposons, until hitting another stop site much further down the array [32]. This way, a single Helitron can engulf the entire 100kb-200kb segment and expand all captured transposons in a concerted manner. If this process is still active in *P. bianor*, a direct test of this hypothesis is to extract and sequence large circular DNA from nuclei, and probe whether the same repeating unit can be recovered with Helitron-specific sites. Since new insertions and deletions can occur between consecutive rolling circle replications, the repeating unit may change over time and contribute to the variety of periodicities among different clusters.

### (b) Similarity between transposon arrays and centromeres

The properties of transposon arrays on Type β chromosomes curiously remind us of centromeres in monocentric organisms. First, centromeres have been shown to be a robust insulator of Hi-C contacts between chromosome arms [6], and transposon arrays have exactly the same effect. This reduction in contact could result from a greater length of Type β chromosomes since most of them have gaps. However, the transposon arrays are gapless on chromosomes 6, 7, 8, and Z, whereas Hi-C contact is still weaker between loci immediately flanking the arrays compared to loci separated by a similar distance on each arm (**Figure 1A**), supporting a mechanism other than length to cause the reduction in contact. Second, meiotic crossovers rarely occur within centromeres [8]. In our study, although it is difficult to estimate local ancestry within transposon arrays, the ancestries of flanking sequences rarely differ, indicating an absence of recombination. Thus, meiotic crossover also appears to be absent within the transposon arrays. Third, centromeres in some insects, such as *Drosophila*, weaken crossover interference between the flanking chromosome arms [33,34], which is exactly what we report here for transposon arrays.

Could transposon arrays be a novel kind of centromere in *P. bianor*? Existing research shows that mitotic kinetochores prefer facultative heterochromatic marks in the silk moth *Bombyx mori* [13]. It is thus reasonable to hypothesize that the likely heterochromatic transposon arrays are also a major region for kinetochore attachment during cell division, which can be tested by antibody staining against conserved kinetochore proteins in metaphase cells. If they do appear to form centromeres, it would imply that Type β chromosomes are monocentric, and that both monocentric and holocentric chromosomes could exist within a single species. If transposon arrays attract more kinetochores in meiosis, it will also be interesting to test if they trigger meiotic drive favoring the transmission of larger arrays, through mechanisms similar to centromere drive [35].

### (c) Why do transposon arrays weaken crossover interference in hybrids?

A caveat in our interpretation of the double crossover data is that the cross was performed between two different species. Without a reference genome for the other grandparent, *P. dehaanii*, it is difficult to attribute all patterns of crossover to features of the *P. bianor* genome.

One of the major mechanistic models for crossover interference is the so-called “coarsening process” [36,37]. In brief, after two homologs pair up (“synapse”) during meiosis, a group of molecules initially distributed throughout the inter-homolog space gradually phase-separate from their solvent via diffusion. The final pool of such molecules will be coarsened (concentrated) at one or more positions along the chromosome, which mark the crossover sites. When coarsening acts efficiently throughout the chromosome, only a single pool exists at the end of the process, preventing secondary crossovers from occurring. This model fits crossover interference patterns well in multiple model organisms [38,39]. Since the molecule can only move through synapsed space between homologs, any disruption to local pairing will be a barrier to the coarsening process and prevent crossover interference from acting across the entire chromosome. Thus, if two hybridizing species diverge strongly at transposon arrays, and homologs fail to pair between the arrays, the coarsening process predicts a decoupling of crossover interference between the flanking “arms”.

Alternatively, crossover interference also decreases with an increasing distance [40,41], similarly to Hi-C contacts as described above. Because most transposon arrays have not been fully assembled, they are likely to be much longer than in the current assembly. The sheer length of the arrays might also be responsible for the weakened crossover interference we observed.

## 4. Conclusion

Tandem repeats are characteristic elements on most monocentric chromosomes in animals and plants. They can selfishly expand at the expense of the host, but some are also essential for chromosome integrity. For monocentric species, centromere function during cell division induces selective forces that influence the abundance and structure of tandem repeats. For holocentric species, however, because the typical constraints on centromeres are absent, it is unclear whether and how selection drives the evolution of tandem repeats. For instance, we do not know if meiotic drive exists on holocentric chromosomes to favor the transmission of larger tandem repeats, nor do we understand how tandem repeats affect the stability of holocentric chromosomes. Our study shows that some Lepidoptera chromosomes can tolerate a significant amount of recently formed tandem repeats enriched for transposons. These repeats appear to affect meiosis by increasing the frequency of double crossovers and the chromosome-wide recombination rate, which might in turn affect their own evolution. Whether these repeats represent a reversion to monocentromeres in holocentric lineages remains an interesting open question.

## 5. Materials and methods

### (a) Genome re-assembly and Hi-C analysis

The source PacBio HiFi reads and Hi-C reads were retrieved from reference [16]. PacBio reads are from one female adult and Hi-C reads are from a different female adult. The mitochondrial genome was first assembled using MitoHiFi [42], and HiFi reads mapped to the mitochondrial genome were filtered out prior to nuclear genome assembly. We tried a variety of parameter combinations with the software hifiasm (0.20.0) [18] (see Supplementary File S1-Table S2), and eventually chose “-D 10 -k 63 -w 51 --max-kocc 1000 --hg-size 520m -s 0.5 -- primary --telo-m CCTAA” as they generated the most contiguous assembly in our trials without producing chimeric contigs between different chromosomes on Hi-C maps. Hi-C reads were then mapped to contigs and scaffolding was performed with YAHS [43]. We manually curated the assembly with Juicebox [44] and ordered chromosomes so that those with transposon arrays were consecutively numbered. We added 13 joins, and added two breaks to remove a single duplicate. To analyze Hi-C contacts, we mapped Hi-C reads to the finished assembly again using the standard Juicer pipeline [44]. Then the output .hic files were converted to a multiresolution .cool file [45] for all downstream analysis using custom scripts (GitHub repository file: 01_Papilio_bianor_HiC_analysis.ipynb). The spectral purity of a Hi-C correlation matrix was computed as one minus the sum of the squared and normalized eigenvalues.

### (b) Repeat annotation

The final assembly was processed with the automated repeat annotation pipeline Earl Grey [46] to construct *de novo* repeat consensus sequences with the options “-c yes -m yes -n 20 -a 5 -e yes -q yes -f 1000”. Using scripts from a manual curation pipeline [47], the raw consensus sequences were blasted against the genome and ranked in a descending order according to the total size of blast hits. We manually curated the most abundant 173 consensus sequences by blasting transposon protein open reading frames or other seed sequences against the genome and extending 3-4kb on each side of matched sequences. All extended matches were aligned and trimmed using visual cues such as the presence of long terminal repeats, terminal inverted repeats, signature terminal motifs, or well-aligned insertion boundaries. We clustered manually curated sequences following an empirical “80-80-80” rule [47] into 116 full-length transposon families. Many Earl Grey consensus sequences were composites of multiple transposons and were split and curated into multiple families. Classification of full-length elements mainly relied on the RepeatClassifier module from the RepeatModeler2 pipeline [48], except for a few sequences which we classified as “Unknown” due to a lack of definitive evidence. To identify satellite DNA, we employed Satellite Repeat Finder [49] and identified eight satellite DNA families with a monomer size greater than 10bp. Most satellite DNA belongs to an abundant 114-bp family. We also removed two rRNA repeats and one histone cluster that were misidentified as transposons from the automated pipeline. All manually curated sequences were combined as a single library for Earl Grey to annotate again, mapping to 46.4% of the genome. We used the script calcDivergenceFromAlign.pl from RepeatMasker [50] to calculate unadjusted Kimura substitution levels between full-length repeat copies.

### (c) Dotplots and higher-order structures

We used ModDotPlot [51] to compute the k-mer-based similarity score between pairs of windows of sizes 1kb, 10kb, and 100kb. The resulting scores were stored as .cool files [45] for downstream analysis. To find major clusters using similarity scores at a resolution of 100kb, the original score matrix was rescaled to 0-1 by the probability density of the scores. The correlation matrix of the rescaled scores was taken as a new measure of similarity, and hence a distance can be defined by subtracting this new similarity score from 1. We used the agglomerative clustering algorithm from the Python scikit-learn package [52] to cluster windows with an average distance smaller than 0.2, an empirical threshold determined by the histogram of all distances (**Figure S5**). The largest 16 clusters are depicted in the main text.

To compute periodicity, we used the dotplot with a resolution of 1kb. Hence, each 1mb sequence defines a sub-dotplot of size 1000x1000. We binarized this sub-dotplot so that only pixels with similarity scores above 99 were taken as 1, and others taken as 0. For each row in this sub-dotplot, we collected the number of pixels (periodicity) between adjacent 1s, and enumerated over all rows. All periodicities from the sub-dotplot were plotted as a histogram with bins taken in log-space. This procedure was repeated for the next sub-dotplot by moving the 1mb windows to the right by a step size of 10kb. The custom scripts for this analysis are in the GitHub repository file “03_Papilio_bianor_Repeat_Visualization.ipynb”.

### (d) Analysis of backcross lines and crossovers

Reads from the three generations of the cross were retrieved from the original paper [22], and mapped to the new assembly, following exactly the same procedure described in that paper using Lep-Map3 [53]. Crossover numbers were estimated by first filtering out likely artifactual breakpoints too close to the telomeres (a known issue with Lep-Map3), then we used Equation 2 from the original paper [22] to estimate crossover numbers, assuming that chromosomes have at most two crossovers per meiosis, a reasonable assumption for butterflies’ short chromosomes. The custom scripts for this analysis are in the GitHub repository file “02_Papilio_bianor_Linkage.ipynb”.

## Acknowledgements

We thank Leah Rosin, Jesper Boman, Xueyan Li, James Mallet, and Jeff Sekelsky for helpful suggestions during the development of this project.

---

**Figure S1.**
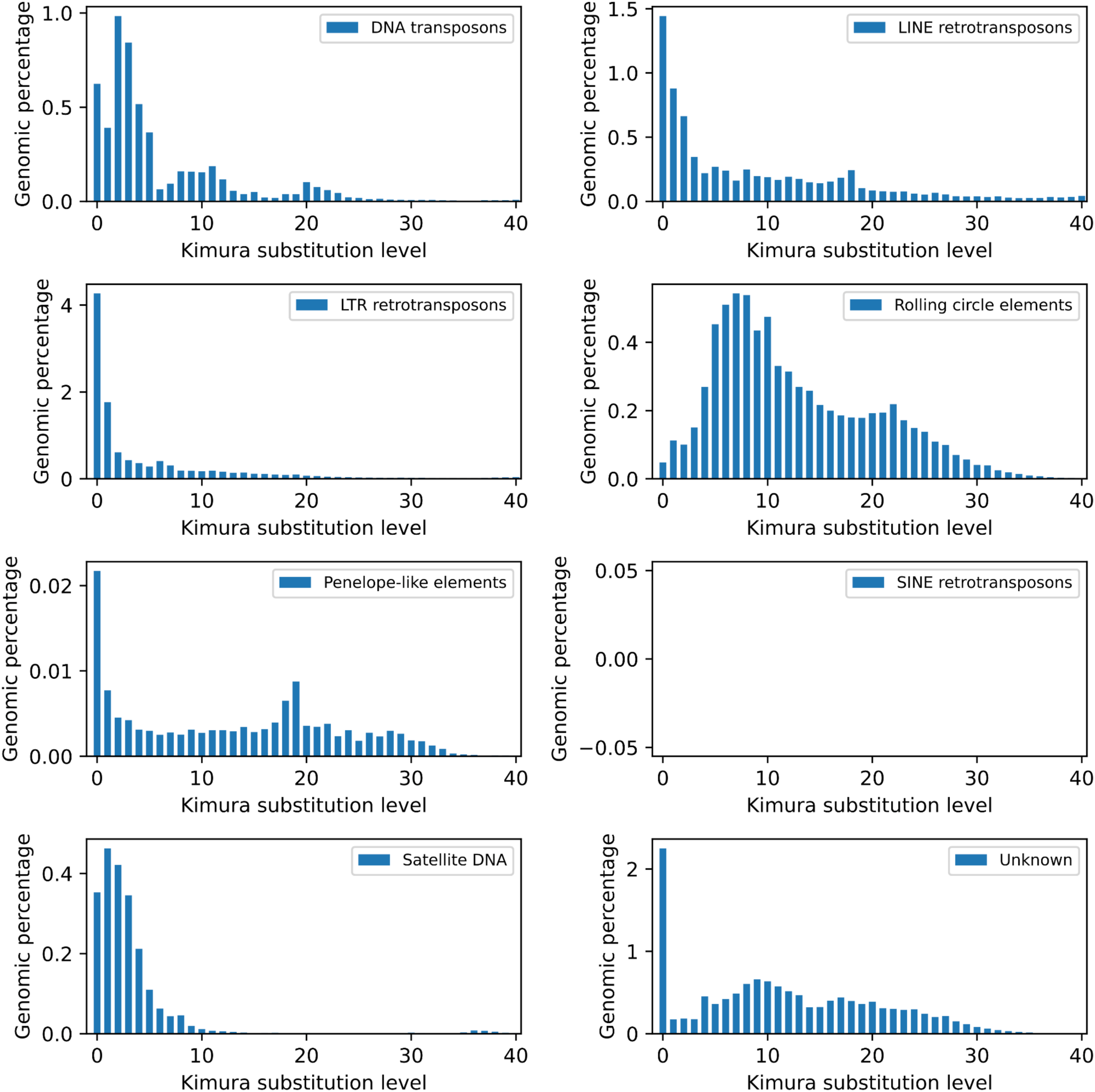
Kimura substitution levels separated by high-level families of manually curated transposons and satellite DNA. We did not identify any SINE elements among the curated families.

**Figure S2.**
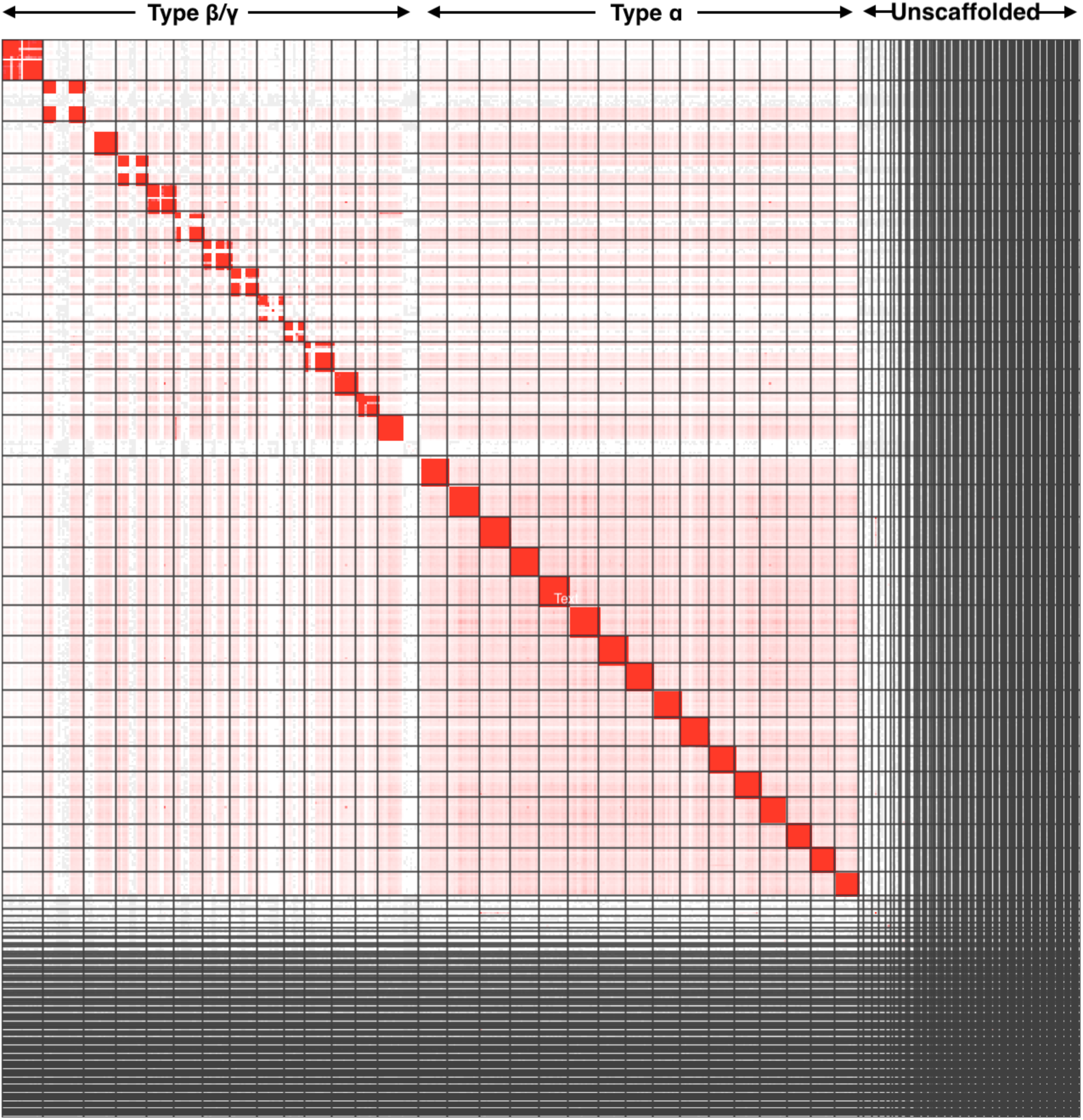
A global Hi-C map of all chromosomes.

**Figure S3.**
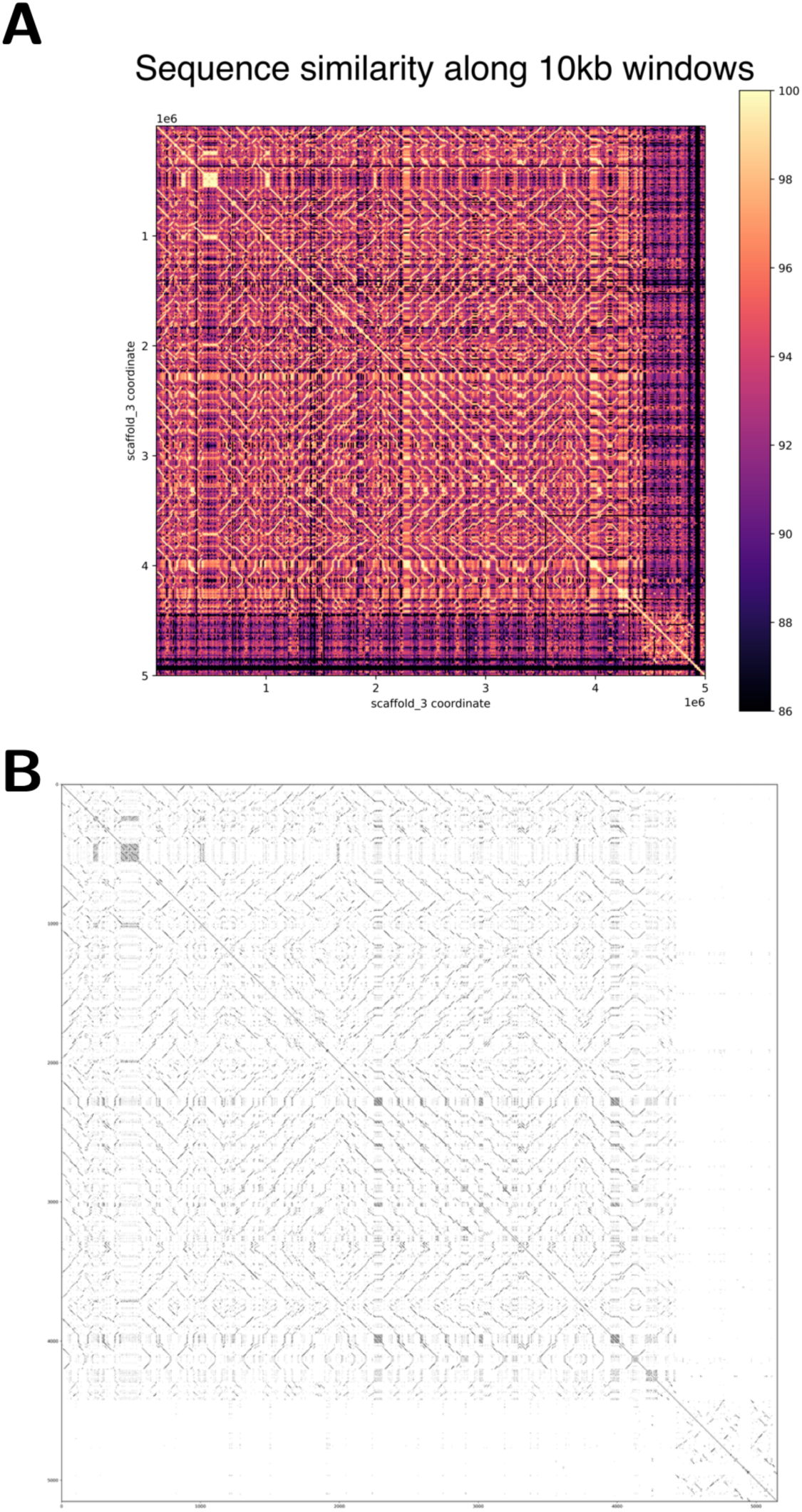
A dotplot of similarity scores across the transposon arrays on Chr3 (cluster 1). **(A)** The dotplot is at 10kb resolution. Both axes are 5mb in length. **(B)** Dotplot of the same region at 1kb resolution, gray-scaled so that highly similar windows are depicted in black.

**Figure S4.**
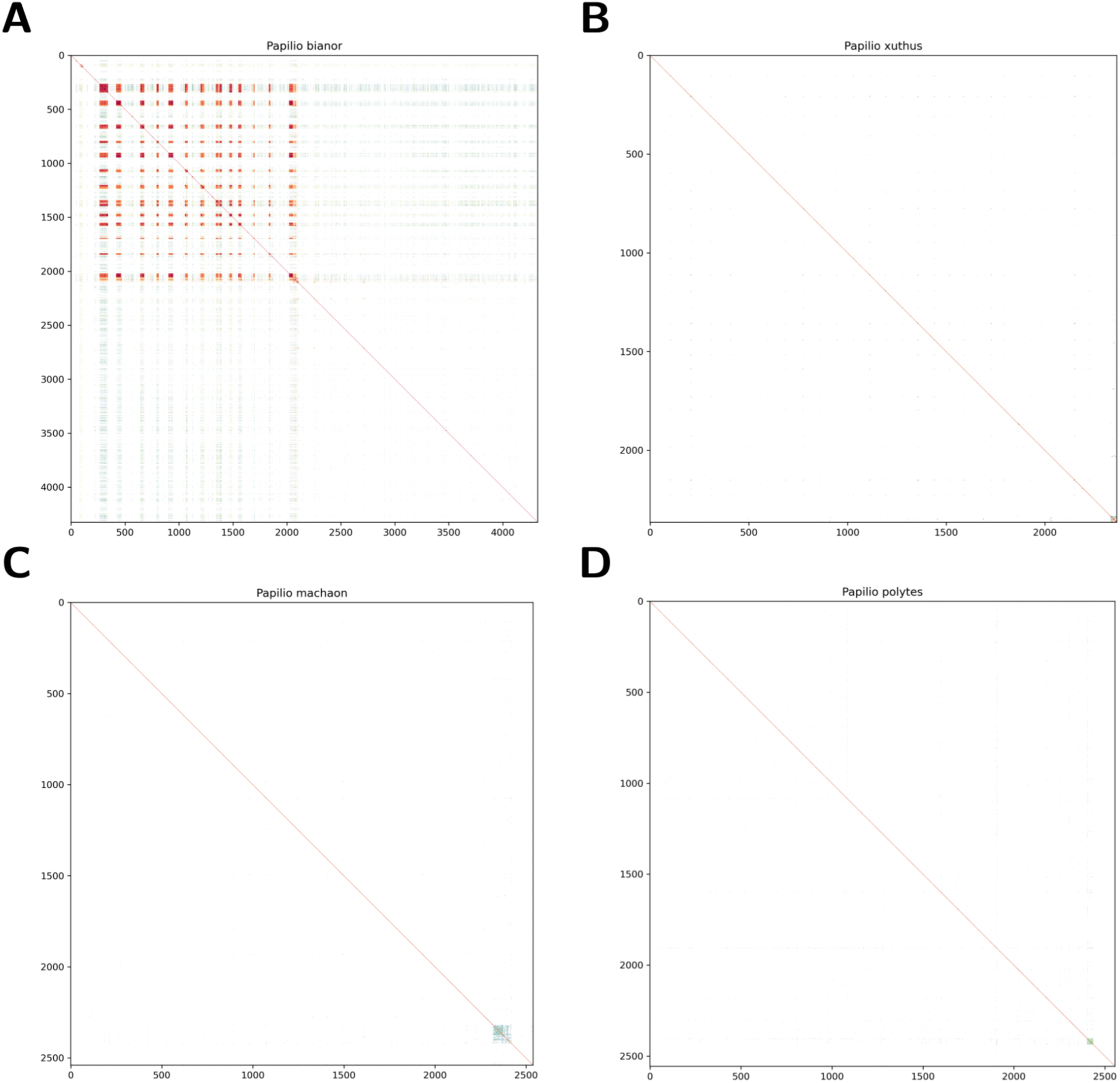
Global dotplots of several *Papilio* genomes with a bin size of 100kb. Highly similar tandem repeats are rare except in *P. bianor*. **(A)** *P. bianor* (this work). **(B)** *P. xuthus*. **(C)** *P. machaon*. **(D)** *P. polytes*.

**Figure S5.**
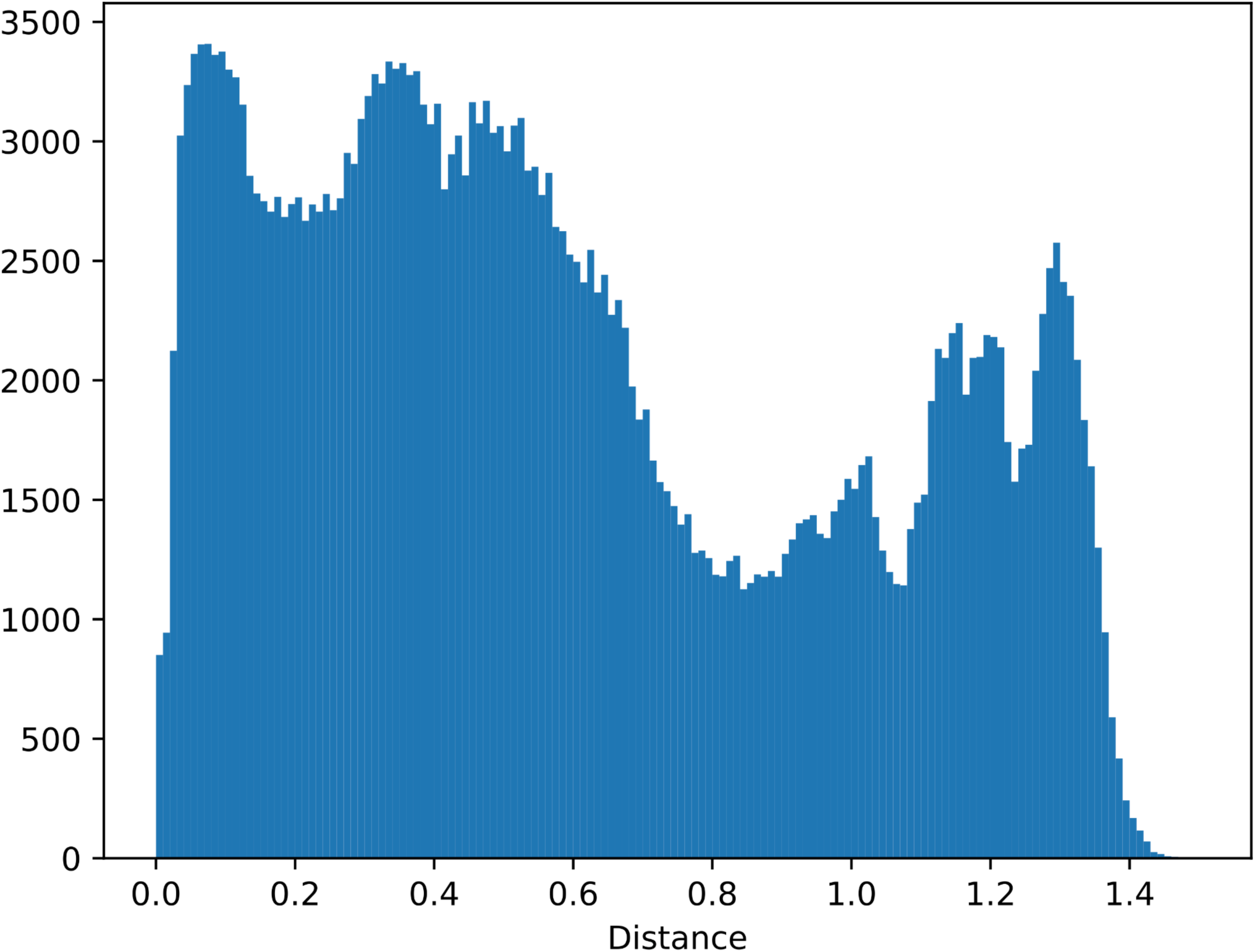
A histogram for re-scaled distances between all pairs of windows in the 100kb-dotplot. The first trough at 0.2 is used as an empirical threshold to delineate clusters.

